# Dynamic Shielding and Allosteric Modulation of Erythropoietin by Glycosylation

**DOI:** 10.1101/2025.08.05.668628

**Authors:** Tomoaki Yagi, Haeri Im, Song-Ho Chong, Yuji Sugita

**Author notes:** Corresponding to Yuji Sugita. **Author Contributions**: T.Y., H.I., and Y.S. designed research; T.Y. and H.I. performed research; Song-Ho Chong contributed new reagents or analytic tools; T.Y. and H.I analyzed data; and T.Y., H.I, and Y.S wrote the paper.

## Abstract

Glycosylation is a ubiquitous and essential post-translational modification that regulates protein structure, solubility, and function. Yet, the mechanisms by which glycans modulate the physicochemical properties of protein surfaces remain incompletely understood. Erythropoietin (EPO), a therapeutic glycoprotein with three N-glycosylation sites, provides a tractable model for dissecting site-specific glycan effects on protein functions. Here, we employ glycan-focused enhanced molecular dynamics simulations, specifically generalized replica exchange with solute tempering (gREST) to overcome the limited sampling of conventional MD and capture the extensive conformational heterogeneity of glycans. Our results demonstrate that glycan effects are highly non-additive: specific combinations of glycosylation sites yield emergent structural outcomes through spatial and dynamical cooperativity. Among them, the N83-linked glycan plays a dominant role in shielding a hydrophobic surface helix, thereby reducing local solvent-accessible hydropathy. Strikingly, the extent of this glycan-mediated surface masking quantitatively correlates with experimentally measured retention times in hydrophobic interaction chromatography, establishing a functional link between molecular-scale shielding and macroscopic behavior. These findings reveal that N-glycans modulate protein surfaces not only through local steric occlusion but also via long-range allosteric effects, providing a new framework for understanding and engineering glycoprotein properties.

## INTRODUCTION

Glycosylation—the attachment of sugar chains to proteins—is one of the most common and vital modifications that occur after proteins are made. It plays essential roles in helping proteins fold correctly, stay soluble but stable, and avoid unwanted immune responses^1^. More than 70% of proteins that are secreted or embedded in cell membranes carry these sugar decorations^2^. The structural diversity of glycans, including variations in composition and branching, enables the precise fine-tuning of protein functions within the cell. Despite their importance, the structural and functional roles of glycans in protein functions remain unclear. Unlike the more predictable protein backbone conformations, glycans are highly flexible, constantly shifting between many conformations^3, 4^. Due to this flexibility, conventional structural approaches make it difficult to visualize the dynamic structures and capture the interaction between glycans and proteins, leaving key questions about their functional roles unanswered.

Erythropoietin (EPO) provides a striking example of how glycosylation governs protein functions^5, 6^. Human EPO is a 30-kDa cytokine with three N-linked glycans (at Asn24, Asn38, and Asn83) and one O-linked glycan, collectively comprising ∼40% of its mass (Fig. 1). This glycan coat is essential for EPO’s in vivo activity and stability: all three N-glycans are required for full biological function, as deglycosylated EPO displays dramatically reduced circulating half-life and potency. Recent advances in glycoprotein chemistry have enabled the production of homogeneous EPO glycoforms with defined glycan patterns, revealing that not only the number but also the position and composition of N-glycans profoundly affect EPO’s pharmacology^7–14^. Notably, glycosylation at Asn83 appears particularly crucial for shielding a hydrophobic patch on EPO’s surface, thereby increasing its solubility and prolonging its residence in the bloodstream. In contrast, glycans at Asn24 and Asn38, while contributing to EPO’s stability, have more modest effects on circulation time and instead may influence folding or receptor engagement. Murakami *et al.* demonstrated that EPO glycoforms bearing a full complement of sialylated N-glycans exhibit the highest erythropoietic activity in vivo, whereas variants missing the Asn83 glycan show accelerated clearance and reduced efficacy^7, 14^. Furthermore, differences in glycan structures (for example, biantennary vs. tetra-antennary sialyloligosaccharides) can subtly alter EPO’s receptor-binding affinity, underscoring that glycan composition as well as occupancy modulates protein–receptor interactions. Together, these findings establish EPO as a paradigmatic glycoprotein whose function is allosterically tuned by its glycan attachments.

**Figure 1.**
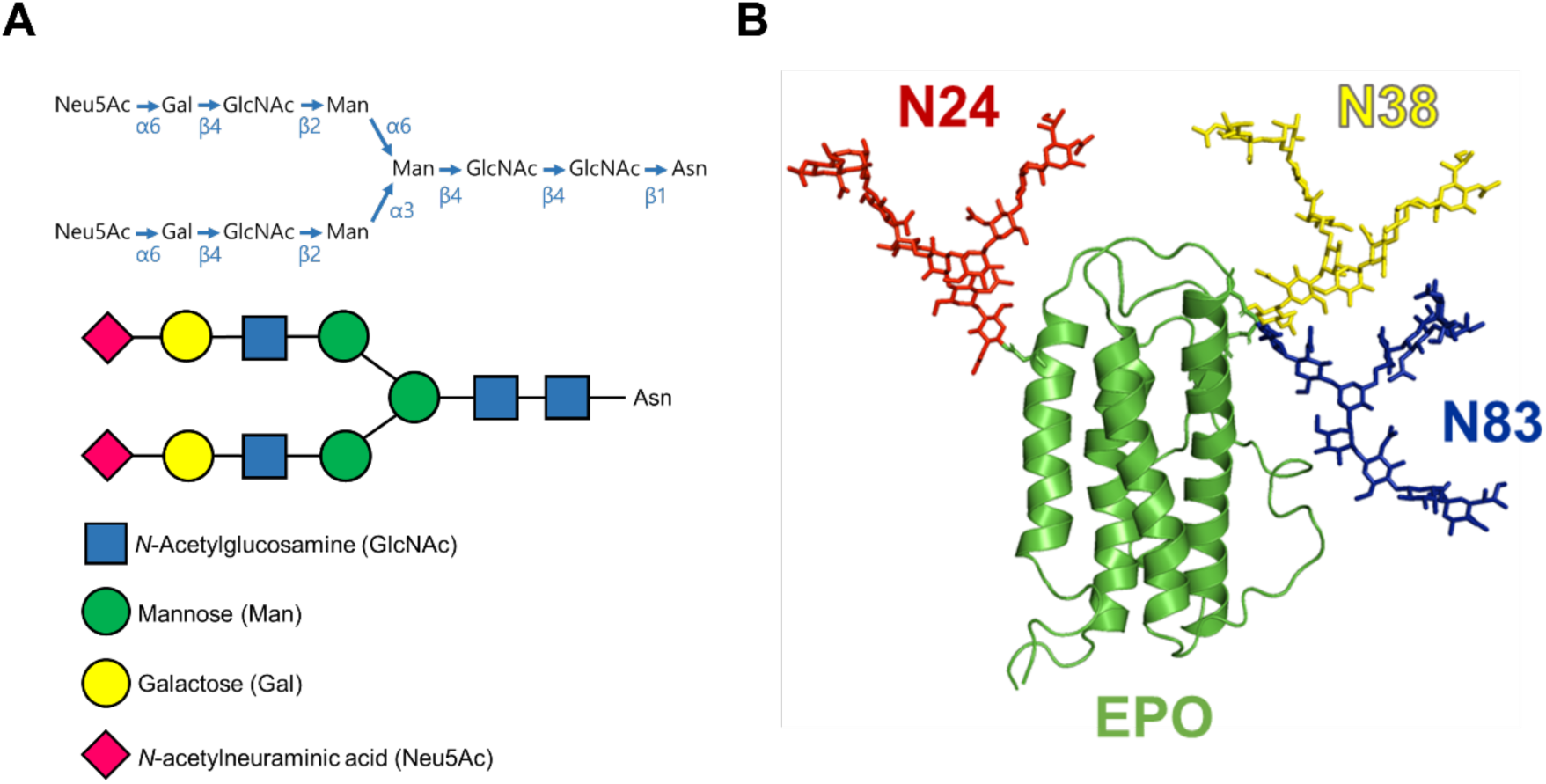
N-glycosylation sites and glycan structural presentation of erythropoietin (EPO). (**A**) Diagram of a biantennary complex-type N-glycan commonly attached to EPO, visualized following the Symbol Nomenclature for Glycans (SNFG). (**B**) Three-dimensional structure of EPO (green) showing N-glycans conjugated at Asn24 (red), Asn38 (yellow), and Asn83 (blue).

Deciphering how glycans exert such effects remains challenging because most structural biology approaches struggle to capture the dynamics of glycans. High-resolution techniques, such as X-ray crystallography and cryo-electron microscopy (cryo-EM), often resolve only the protein backbone and sidechain^4^. At the same time, glycans appear as blurred electron density or are absent due to disorder and heterogeneity. Even when glycans are present in crystal structures, they typically terminate at the first one or two saccharide units, leaving the extended glycan conformations unresolved. Similarly, computational structure prediction has limits: AlphaFold^15, 16^ and related AI-based models, while revolutionary for globular protein domains, generally cannot account for post-translational modifications and tend to misrepresent highly flexible regions. For heavily glycosylated proteins, a single static model is inadequate – glycan-coated proteins, such as EPO, exist as dynamic ensembles that static approaches cannot easily portray. The net result is a gap in our structural and mechanistic understanding: experimental snapshots give only a partial view of glycoprotein surfaces, and thus the specific ways that glycans mask, stabilize, or rearrange protein epitopes remain difficult to discern.

Molecular dynamics (MD) simulations have emerged as a powerful complementary approach to probe the behavior of glycoproteins in atomistic detail. By explicitly simulating thermal fluctuations, solvation, and glycan movements, MD can capture the dynamic shielding and allosteric influences that glycans impart – phenomena often invisible to static methods. Over the past decade, simulations have provided key insights into glycan functions across various systems. In the case of HIV Env, enhanced MD simulations revealed an extensive network of inter-glycan contacts and transient openings in the glycan shield, rationalizing how certain antibody epitopes become accessible^17, 18^. Similarly, simulations of the SARS-CoV-2 spike have illustrated how specific N-glycans modulate the protein’s breathing motions and receptor-binding domain accessibility, effectively gating the transition between receptor-inactive and active states^19–22^. Beyond viral proteins, MD has been applied to human glycoproteins (antibodies, receptors, etc.), demonstrating that glycans can restrain flexible loops, stabilize particular conformational states, or sterically hinder protein-protein interactions. These computational studies underscore that glycans are not mere decorations but dynamic structural elements: by continually moving and rearranging, they can shield large swathes of a protein’s surface or subtly bias the protein’s conformational ensemble. MD simulations thus provide a window into these glycan-mediated effects, offering mechanistic hypotheses that complement experimental observations.

At the same time, accurately simulating glycoproteins presents its own challenges. The flexible, high-dimensional conformational space of glycans – with numerous rotatable bonds in each oligosaccharide chain – results in a rugged free-energy landscape characterized by multiple minima and substantial barriers between them. Achieving sufficient sampling of glycan conformations (especially for complex branched N-glycans) can be prohibitively slow with conventional MD alone, which may become trapped in one region of phase space. To overcome this sampling problem, enhanced-sampling techniques have been increasingly applied to glycoproteins. In particular, replica-exchange methods – which involve simulating multiple copies (replicas) of the system under different conditions and periodically swapping them – have proven effective at overcoming energy barriers in glycan rotation. Temperature replica-exchange MD (REMD)^23–26^ and Hamiltonian replica-exchange (HREX)^27, 28^ methods enable glycan torsions to escape local minima by visiting higher-temperature or biasing Hamiltonians. The conformational ensembles are available by collecting the trajectories of the replicas. Recent developments go even further: for example, Yang *et al.* combined replica exchange with selective biasing of glycan dihedral angles (a method termed HREST-BP)^18^ to capture the full conformational heterogeneity of the HIV Env glycans, revealing many glycan orientations that were inaccessible in shorter simulations. Likewise, Grothaus *et al.* reported a hybrid replica-state exchange scheme that couples REST with collective-variable tempering (REST-RECT)^29^, enabling transitions over all relevant torsional barriers and yielding converged ensembles for complex N-glycans in solution^30^. The upshot is that we can now begin to fully explore the dynamic repertoires of glycans and their impact on protein surfaces, rather than viewing glycans as static modifiers. In this study, we utilize generalized replica exchange with solute tempering (gREST), a replica-exchange method that selectively enhances the sampling of predefined solute regions. By restricting the solute tempering region to N-glycans alone, gREST allows exhaustive exploration of glycan conformational space without perturbing the underlying protein structure. This targeted acceleration enables us to capture the full dynamic repertoire of glycans while maintaining the native-like conformational ensemble of the protein. By combining the enhanced-sampling MD with experimental insights, we aim to elucidate a dynamic model of glycosylation’s influence on EPO – one in which glycans act as movable shields and allosteric effectors, rather than passive decorations, thereby paving the way toward a deeper understanding of glycoprotein function and design.

## RESULTS

### Glycan Conformational Sampling Efficiency by gREST

We investigated the conformational dynamics of seven EPO proteins with different N-linked glycosylation at Asn24, Asn38, and Asn83 in explicit water solution. Three systems contain one glycosylation (N24, N38, and N83). At the same time, the other four include multiple glycosylations (N24N38, N24N83, N38N83, and N24N38N83). For comparison, the structure and dynamics of EPO protein without any glycosylation (epo) was also investigated. To quantify the conformational sampling efficiency using enhanced sampling algorithms, we conducted atomistic MD simulations based on generalized replica exchange with solute tempering (gREST)^31^ and conventional MD (CMD) simulations. After exploring multiple glycosylation states of the target protein, the glycan conformations were subjected to k-means clustering (k = 2), using the Cartesian coordinates of heavy atoms for each glycan moiety extracted from the trajectories as input. The temporal evolution of the cumulative cluster populations was then analyzed to assess convergence behavior (Fig. 2). To visualize the conformational variability within each cluster, 50 frames were randomly selected from each cluster and structurally superimposed to generate representative structures, as shown.

**Figure 2.**
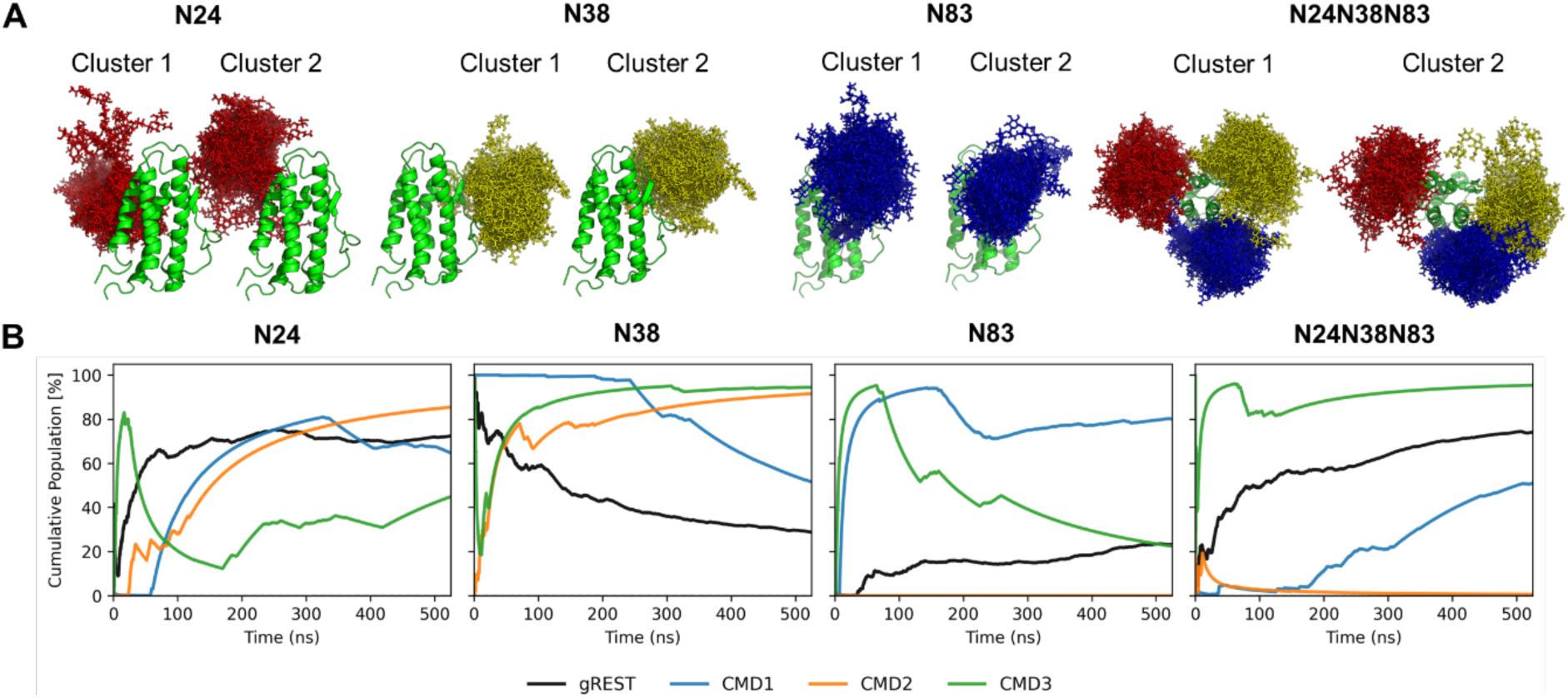
Assessment of glycan conformational sampling using gREST and conventional MD simulations. (**A**) Representative glycan conformations sampled from each of the two major clusters identified by k-means clustering (k = 2), performed on the Cartesian coordinates of glycan heavy atoms. For each cluster, 50 frames were randomly selected and superimposed to illustrate structural diversity within the cluster. (**B**) Time evolution of the cumulative population of each cluster for different glycosylation sites. Each line denotes a single simulation: gREST (black) and three conventional MD trajectories (CMD1–3; blue, orange, green). In singly glycosylated systems, the clusters correspond to protein-bound and solvent-exposed glycan states, while in multiply glycosylated systems, they represent configurations with and without inter-glycan contacts—most notably between Asn38- and Asn83-linked glycans. The results highlight the enhanced sampling efficiency of gREST over conventional MD.

For singly glycosylated systems, the two identified clusters predominantly correspond to glycan conformations engaged in protein surface interactions and those exhibiting extended solvent exposure. In systems with multiple glycans, clustering distinguished between compact conformations featuring inter-glycan contacts (notably between N38- and N83-linked glycans) and spatially segregated glycan states. CMD trajectories showed slow and often incomplete population convergence within 500 ns, reflecting limited exploration of the glycan conformational landscapes. In contrast, gREST simulations achieved more rapid and thorough sampling of both clusters, highlighting its superior efficiency in overcoming the entropic and enthalpic barriers that characterize glycan dynamics. Hereafter, we mainly focus only on the gREST trajectories at room temperatures to investigate the glycan dynamics, its interaction with EPO protein, and glycan-glycan interactions.

### N-Glycosylation Modulates Local Flexibility and Global Conformational Dynamics of the Protein

To evaluate how N-glycosylation influences protein conformational dynamics, we computed the residue-wise root-mean-square fluctuations (RMSF) of Cα atoms across various glycosylation states. The differential RMSF (ΔRMSF), defined as the difference between glycosylated and non-glycosylated systems, is shown in Figure 3A. Notably, glycan attachment not only alters the flexibility of residues in close proximity to the glycosylation sites—N24, N38, and N83—but also induces pronounced changes in regions distal to the modification sites. In particular, several loop regions located far from the glycosylation sites exhibit increased or suppressed fluctuations depending on the glycan composition. For example, enhanced flexibility is observed around residues 110–140, despite these regions being spatially distant from N24 and N38. This indicates that N-glycosylation can exert long-range allosteric effects, modulating the dynamics of remote structural elements such as loops involved in inter-helical packing or functional interfaces. These observations underscore the role of glycans as modulators of not just local but also global dynamic behavior in glycoproteins.

**Figure 3.**
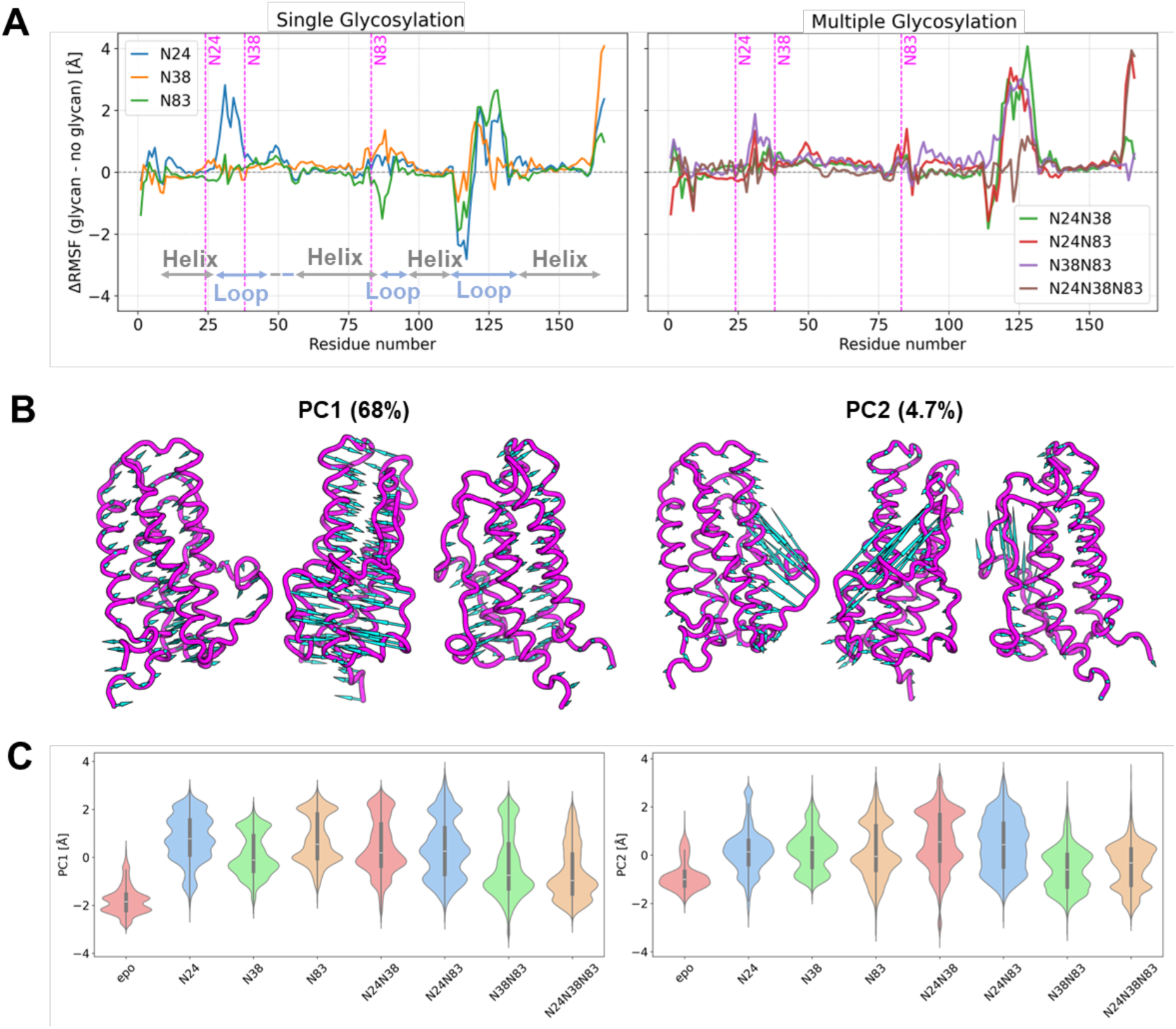
Effects of N-glycosylation on protein conformational fluctuations and collective motions. (**A**) Residue-wise difference in root-mean-square fluctuation (ΔRMSF) of Cα atoms between glycosylated and non-glycosylated EPO for single (left) and multiple (right) glycosylation states. Positive values indicate increased flexibility upon glycosylation. Loop and helix regions are annotated based on secondary structure assignment. (**B**) Principal component vectors mapped onto the PDB protein structure for visual comparison. Magenta ribbons represent the reference structure; cyan arrows depict the direction and amplitude of PC1 and PC2 motions. (**C**) Distribution of principal component (PC1 and PC2) projections derived from PCA on the Cα Cartesian coordinates of the protein for all glycosylation states. PC1 represents collective expansion/contraction motion, whereas PC2 corresponds to loop dynamics (residues 112–137). Glycosylation leads to substantial shifts in the distributions relative to the non-glycosylated form.

To gain mechanistic insight into the nature of the collective motions affected by glycosylation, we visualized the first (PC1) and second (PC2) principal components obtained from PCA^32^ of the Cα Cartesian coordinates by mapping the corresponding eigenvectors onto the protein structure (Figure 3B). These modes reveal two distinct classes of motion, each differentially modulated by the presence or absence of N-glycans. PC1 corresponds to a global expansion–contraction motion of the protein scaffold. In the non-glycosylated system, this mode exhibits relatively small amplitude, indicating limited structural breathing. In contrast, glycosylated systems show a marked increase in PC1 amplitude and a shift of the distribution toward positive values (Figure 3C left), indicating a bias toward expansive conformational states. This suggests that glycosylation increases the dynamic range of the protein, favoring more open structures. Notably, the degree of this shift depends on the glycosylation pattern, with specific combinations (e.g., N24N38) inducing stronger expansion than others. PC2 represents localized motion of the loop region spanning residues 112–137. In the absence of glycans, the distribution of PC2 values is centered near zero, reflecting symmetric fluctuations of the loop. Upon glycosylation, the distribution shifts strongly in the positive direction, indicating a preferential movement of the loop toward the helical core. This motion corresponds to a partial enclosure of the helices by the loop, effectively reshaping the protein surface. The shift in PC2 is most pronounced in the N38N83 and N24N38N83 glycoforms, correlating with enhanced loop fluctuation in RMSF profiles.

These observations demonstrate that N-glycosylation induces systematic and site-specific modulation of both global and local protein dynamics. In particular, glycosylation at EPO not only expands the protein conformation but also directs loop motion toward more compact and shielded states—effects that are absent or minimal in the non-glycosylated form. This highlights the critical role of glycans in sculpting the conformational ensemble of glycoproteins.

### Cooperative Spatial Organization of Glycans on the Protein Surface

To elucidate how glycosylation modulates the spatial organization of glycans relative to the protein and to each other, we conducted a geometric analysis of both glycan–protein and glycan–glycan interactions. We defined an angle for each glycosylation site as the angle formed by the vector from the Cα atom of the Asn residue to the protein center of mass, and the vector from the Asn to the glycan center of mass. This angle serves as a proxy for glycan proximity to the protein surface—smaller angles indicate closer contact. Additionally, we measured the pairwise distances between glycan centers of mass to evaluate inter-glycan interactions.

The angle distributions (Figure 4C right) indicate that the glycan at N83 consistently adopts conformations closer to the protein surface, exhibiting the smallest median angle across all glycosylation states. N24 and N38 also exhibit surface proximity but with broader distributions and higher average angles, suggesting more conformational variability. These observations are in line with the representative cluster structures obtained via k-means clustering and their respective populations. Interestingly, increasing the number of glycans leads to a systematic shift of the angle distributions toward larger values for all sites. This suggests that inter-glycan crowding or repulsive interactions push the glycans away from the protein surface. This shift is particularly pronounced in the N38 and N83 angles, which show significant broadening in the triple glycosylation state. These trends underscore the context-dependent, cooperative nature of glycan–protein positioning.

**Figure 4.**
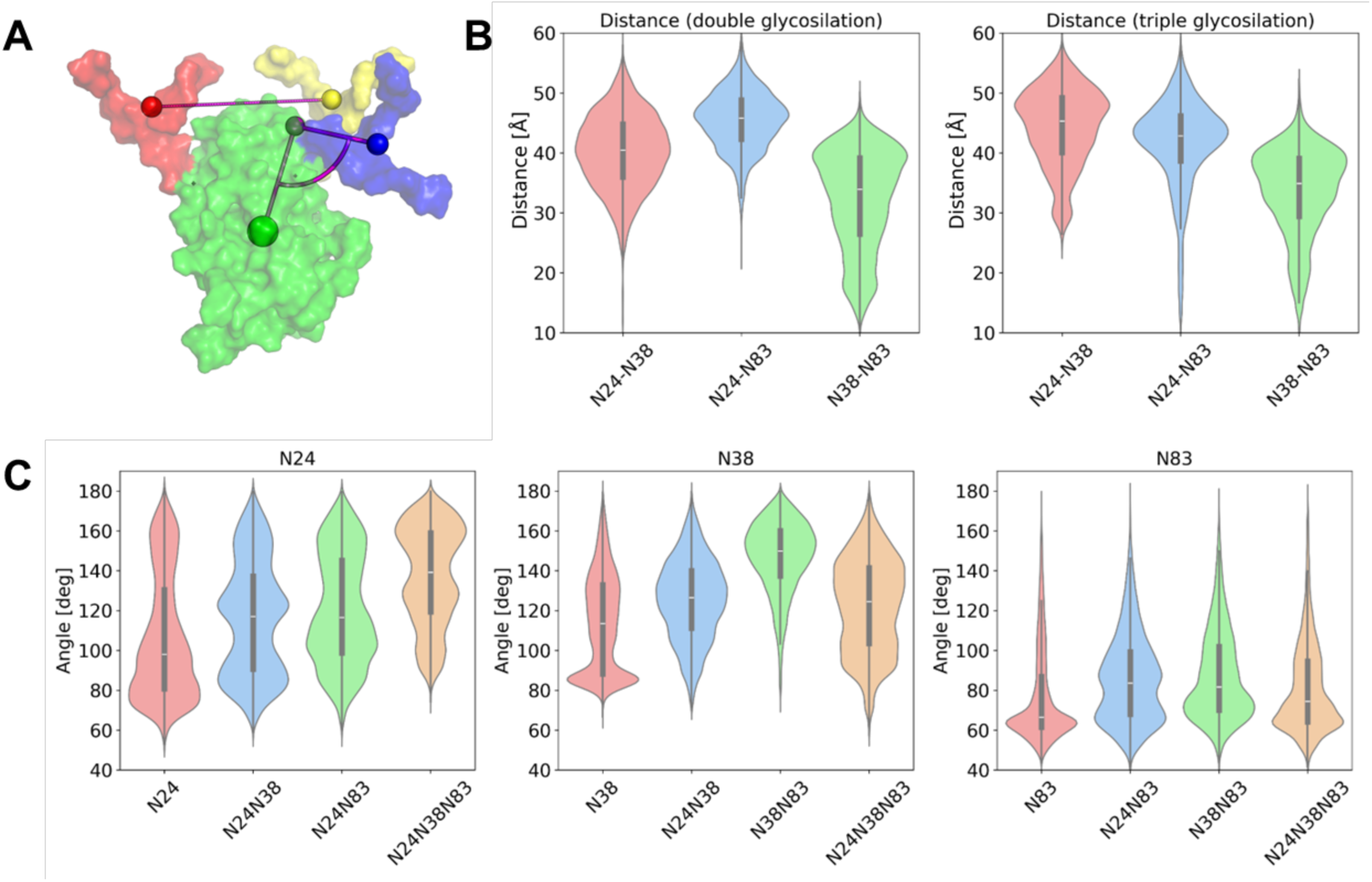
Distributions of glycan–protein angles and inter-glycan distances across EPO glycoforms. (A) Structural illustration of erythropoietin (EPO) showing the geometric definitions of inter-glycan distance and angle. Each glycosylation site is represented by a colored sphere at the glycan’s center of mass (N24: red, N38: yellow, N83: blue), and lines connecting them denote the measured distances. Angles are defined at each glycosylation site as the angle formed between vectors connecting the other two glycans. (B) Distributions of the inter-glycan center-of-mass distances (in Å) between each glycan pair (N24–N38, N24–N83, N38–N83), shown for double (left) and triple (right) glycosylated states. (C): Distributions of inter-glycan angles (in degrees), defined at each glycosylation site (N24, N38, N83) across different glycosylation states.

Pairwise distances between glycan centers of mass (Figure 4B) reveal that the N38–N83 pair tends to approach most closely, indicating a preferred interaction interface between these two glycosylation sites. In contrast, N24–N83 and N24–N38 pairs exhibit larger typical distances. In triple glycosylation states, all glycan–glycan distance distributions become broader and shift toward longer values. This observation suggests a tug-of-war effect, in which the formation of close contact between one glycan pair (e.g., N38–N83) forces the remaining glycans into more extended conformations due to spatial competition. This long-range balancing behavior reflects the constrained geometry of the glycoprotein surface and highlights the non-additive, competitive nature of multivalent glycosylation.

### Site-Specific Glycan–Protein Interactions Mediated by Electrostatics and Surface Hydrophobicity

To investigate the spatial characteristics of glycan–protein interactions, we computed residue-wise contact frequencies for each glycosylation site (Figure 5, left). Contacts were defined as instances where any heavy atom of the glycan was within a specified distance threshold of a protein residue. The resulting profiles reveal that glycan–protein interactions occur primarily near the glycosylation sites, as expected. However, the interaction patterns differ across sites. The glycan at N83 shows sharp contact peaks in residues immediately adjacent to the glycosylation site, consistent with its deeply embedded orientation suggested by angle distribution. The contacts are localized predominantly on α-helices, particularly the region spanning residues 55–85. In contrast, N24 and N38 also interact with the protein surface but with broader and more variable contact distributions. Notably, N38 engages not only helices but also loop regions, reflecting the smaller flexibility of the loop as show in the RMSF (Fig 2).

**Fig. 5.**
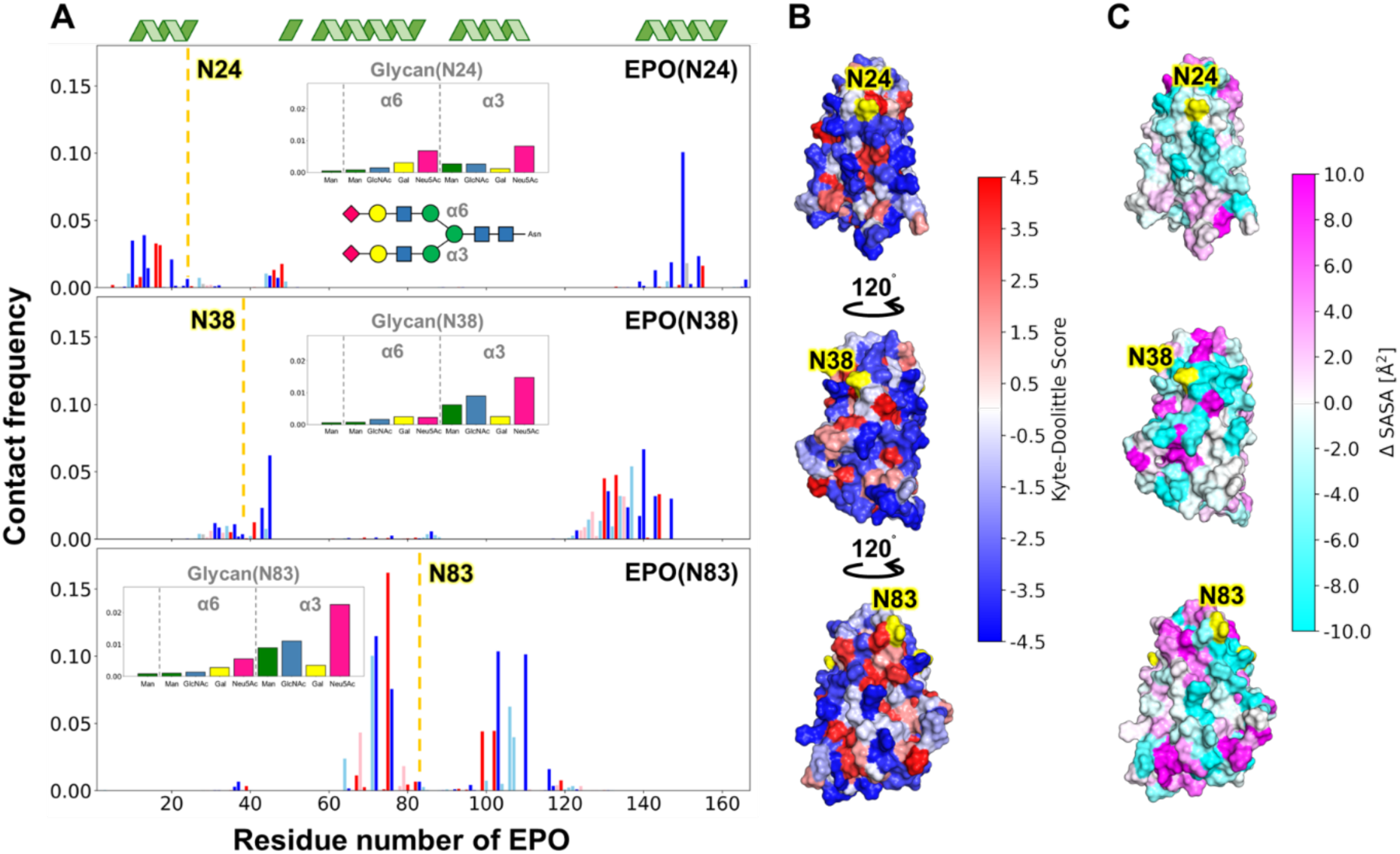
Contact frequency between EPO protein residues and N-linked glycans at N24, N38, and N83. (A) Residue-wise contact frequency between each glycan and the protein for N24, N38, and N83 glycosylation sites. Contact is defined by proximity between heavy atoms, with blue and red bars corresponding to α6- and α3-linked glycans, respectively. Insets show the glycan composition and linkage types. (B) Molecular surface of the protein colored by residue hydrophobicity score, highlighting the spatial distribution of hydrophobic (red) and hydrophilic (blue) residues. Notably, hydrophobic residues are enriched near the N83 site, especially in the α-helix spanning residues 53–86. (C) Difference in solvent-accessible surface area (SASA) between glycosylated and non-glycosylated states mapped onto the protein surface. Glycosylation leads to reduced SASA near the attachment sites, with the largest decrease observed at N83, particularly in the hydrophobic helical region.

Most frequent contacts arise from the terminal moieties of the glycans, particularly the α2,3- or α2,6-linked sialic acids, which carry negative charges. These negatively charged sugar units tend to interact preferentially with positively charged surface residues such as Lys and Arg, suggesting an electrostatically driven interaction mechanism. This observation is supported by the high contact frequencies localized at basic residues in the proximity of each glycan’s terminal branch.

To evaluate the relationship between contact patterns and residue properties, we mapped the residue hydrophobicity score onto the protein surface (Figure 5B). Around the N83 site, strong contact regions coincide with clusters of hydrophobic residues, especially in the α-helix spanning residues 55–85. In contrast, the N24 and N38 sites are located in more hydrophilic environments, which correlates with fewer persistent contacts and more dynamic glycan conformations. This suggests that local surface hydrophobicity contributes to stabilizing glycan–protein interactions, especially at N83.

To further understand how glycosylation affects surface exposure, we calculated the difference in solvent-accessible surface area (SASA) between glycosylated and non-glycosylated forms, and mapped the results onto the protein structure (Figure 5C). Consistent with the contact analysis, glycosylation leads to local reductions in SASA around each glycosylation site. The most pronounced decrease occurs at N83, particularly within the hydrophobic α-helical region, indicating that the glycan at this site effectively shields hydrophobic residues from solvent exposure. This shielding effect is less prominent at N24 and N38, where the glycans are more solvent-exposed and flexible.

### Hydrophobic Helix Near N83 Controls Retention Behavior via Surface Accessibility

To investigate how glycosylation modulates the interaction between EPO and the hydrophobic surface of the chromatography column, we analyzed the relationship between the solvent-accessible surface area (SASA) of EPO and its retention time in high-performance liquid chromatography (HPLC) obtained by Murakami et al.^7^. Since retention time in this context reflects the strength of hydrophobic interactions with the column matrix, we used SASA as a structural proxy to infer which regions of the protein contribute to this behavior.

We computed both total and hydrophobic SASA values for various residue subsets, including the entire protein (residues 1–166), the hydrophobic helix (residues 55–85), and extended regions surrounding the helix (residues 55–115 and 55–130). Correlation analysis (Figure 6B) revealed that the hydrophobic helix near the N83 site shows the strongest positive correlation with retention time (R = 0.96 for total SASA). This suggests that increased surface exposure of this helix enhances EPO’s retention on the hydrophobic column. By contrast, no correlation was observed between retention time and the total SASA of the entire protein (R = –0.23), underscoring the localized nature of the interaction. When extended regions including adjacent helices and loops were considered, the correlation weakened (R = 0.87 and 0.89), further supporting that the core hydrophobic helix itself is the dominant contributor to the observed chromatographic behavior.

**Fig. 6.**
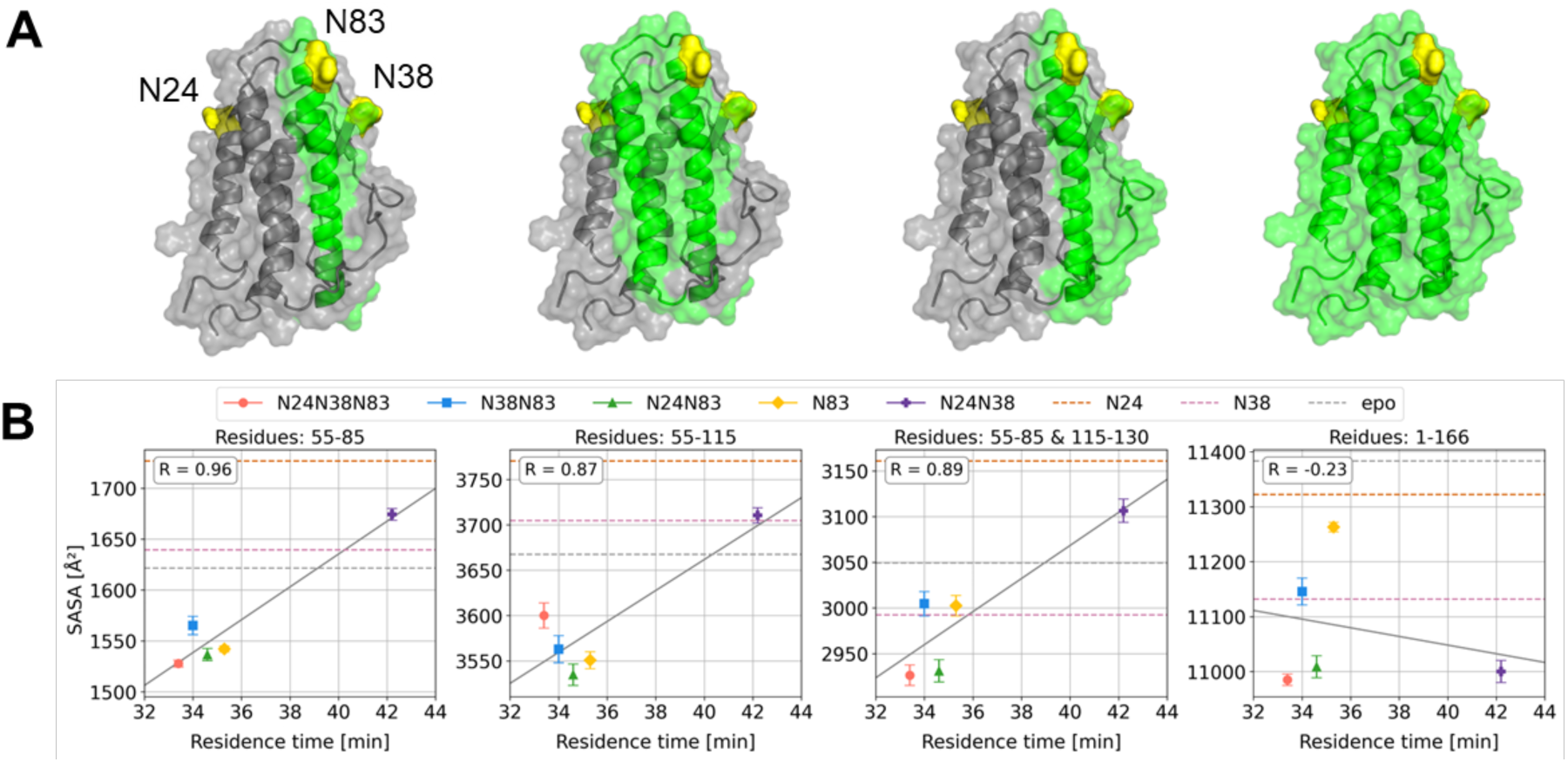
Glycosylation-dependent shielding of hydrophobic helices determines HPLC retention. (A) Molecular surfaces of representative EPO glycoforms expressing the selected SASA area in green and the residual area in gray and N-glycosylation sites (yellow). While multi-glycosylated species such as N24N38N83 exhibit extensive shielding of hydrophobic patches, glycoforms like N24N38 expose large hydrophobic surfaces. (B) Correlation between HPLC retention time and solvent-accessible surface area (SASA) for different residue groups. Each point represents a distinct glycoform; dashed lines indicate values for N24, N38, and wild-type glycosylated EPO. SASA of helix 55–85 showed the strongest correlation with retention time (R = 0.96), consistent with the observed retention order: N24N38N83 < N38N83 < N24N83 < N83 << N24N38. Whole-surface SASA (residues 1–166) showed no clear relationship with retention (R = −0.34), indicating that local hydrophobic exposure, rather than global surface area, is the primary determinant of retention in this system.

The surface renderings (Figure 6A) show that glycan positioning—particularly at N83—can directly affect the exposure of the hydrophobic helix. In glycoforms with bulky glycans near N83, the helix becomes more buried, reducing SASA and shortening retention time. These results support the hypothesis that glycan-mediated masking of the hydrophobic helix modulates retention behavior by altering hydrophobic surface accessibility.

Notably, the N38N83 glycoform deviates from the regression line for the SASA of the hydrophobic α-helices. This outlier likely reflects glycan-specific effects that are not fully captured by a scalar SASA metric of the helix—e.g., cooperative packing of the N38 and N83 glycans that alters EPO’s orientation at the column surface, transient shielding due to glycan–helix motions, or microheterogeneity in the glycans that changes local hydrophilicity. These nonlocal and dynamic factors can weaken the otherwise tight link between helix accessibility and retention.

## DISCUSSION

Our study reveals that *N*-glycans on erythropoietin (EPO) are not passive structural appendages but dynamic modulators that reshape protein surfaces, alter conformational ensembles, and directly influence pharmacological behavior. By employing generalized replica exchange with solute tempering (gREST), we captured the conformational heterogeneity and cooperative organization of EPO’s glycans, phenomena largely invisible to classical structural approaches such as X-ray crystallography or cryo-EM, which often truncate or omit glycans due to their inherent flexibility. Recent computational frameworks, including GlycoSHAPE^33^ and GlycoSHIELD^34^, have significantly advanced the modeling of glycoprotein surfaces by grafting pre-sampled glycan ensembles or estimating shielding effects from static structures. These methods provide rapid and practical means for reconstructing glycosylation patterns, especially when experimental density maps are incomplete. However, they rely on pre-computed conformer libraries and thus cannot fully account for the coupled, cooperative motions of glycans and protein backbones. In contrast, our gREST-MD simulations explicitly capture such dynamics, revealing features such as inter-glycan steric competition and allosteric modulation of distal loops. We consider these approaches complementary rather than competing: rapid conformer-based modeling (e.g., GlycoSHAPE or GlycoSHIELD) can identify candidate glycan orientations or shielding zones, while enhanced-sampling MD provides a mechanistic, time-resolved picture of glycan–protein interactions.

### N83: the dominant surface-hydropathy modulator

Among the three N-glycosylation sites on EPO, N83 plays the most dominant and multifaceted role in controlling both structure-exposure relationships and pharmacologically relevant properties. Experimental studies using homogeneous glycoforms provide compelling evidence: even a single glycan at N83 is sufficient to decrease RP-HPLC retention time, indicating reduced surface hydrophobicity, while N24/N38 double glycosylation, lacking N83, produces the most hydrophobic and aggregation-prone species, despite a greater total glycan count^7^. These results highlight that positional identity surpasses glycan number in determining surface properties.

Our simulations explain this through a local, time-resolved shielding mechanism: the N83-linked glycan frequently adopts compact, surface-associated conformations, maintaining close contact with a hydrophobic α-helix (residues 55–85) that is solvent-exposed in the non-glycosylated form (Fig. 5). This glycan–helix association significantly reduces the local SASA, and the extent of this reduction correlates strongly with HIC retention time (Fig. 6), establishing a direct, mechanistic link between molecular-scale dynamics and macroscopic elution behavior.

Structurally, the region shielded by N83 is enriched in nonpolar residues (Leu, Ile, Val, Phe) and flanked by basic residues (Lys, Arg), creating a favorable environment for hydrophobic interactions and electrostatic stabilization with the negatively charged sialylated termini of the glycan. This mixed-mode interaction (steric, hydrophobic, and electrostatic) supports persistent glycan–protein contact over the course of simulation trajectories, yet remains dynamically flexible—a mobile shield, not a static blocker.

Biologically, this dynamic masking provides a dual benefit: it prevents premature aggregation or ER retention, as evidenced by lower glycosylation signals in UGGT recognition assays for N83-containing glycoforms^11^, and it prolongs serum half-life, as N83 helps reduce renal clearance by suppressing non-specific interactions with hydrophobic interfaces.

Importantly, the shielding effect of N83 is not binary but concentration- and structure-dependent. For example, replacing a tetra-antennary, fully sialylated glycan with a bi-antennary or a sialylated form is expected to weaken coverage and increase HIC retention and aggregation risk. Our simulations are well-positioned to capture these subtleties, offering a framework to quantitatively predict how specific glycan architectures at N83 tune shielding efficiency.

### N38: a cooperative modulator and kinetic damper

Experimental studies using structurally uniform EPO glycoforms show that N38, especially when paired with N83, plays a more important functional role than N24. Among two-site glycoforms, those that include N38 and N83 show higher biological activity than those with N24 and N83, highlighting a site-specific hierarchy in functional importance^7^. Beyond activity, N38 also appears to support protein folding: removing N38 leads to lower oxidative folding yields and more misfolded protein products. For example, when folding was conducted at low concentration (0.01 mg/mL), the glycoform containing N38 and N83 achieved ∼90% correctly folded protein, while N24/N83 and N83-only variants yielded only ∼66% and ∼63%, respectively^7^.

In our simulations, the N38 glycan does not act alone—it dynamically interacts with the N83 glycan, alternating between compact conformations that cooperate to shield the hydrophobic surface, and extended conformations that reduce steric clashes. This dynamic behavior helps to stabilize the extent of surface exposure over time, not just on average, but by reducing fluctuations (i.e., lowering variance). These insights match experimental results showing that although N24N38 and N38N83 glycoforms contain the same number of glycans, the latter has significantly lower hydrophobicity in RP-HPLC (33.99 vs 38.64 min retention)^7^, suggesting better surface masking when N38 partners with N83.

### N24: a hydrophilic additive, not a core modulator

In contrast to N38 and N83, the N24 site contributes least to EPO’s biological activity in vivo. Retention times in RP-HPLC further confirm that N24N38 (no N83) is the most hydrophobic glycoform tested, whereas a single N83 glycan produces less hydrophobic retention than N24N38, despite having fewer total glycans. This indicates that the location of a glycan is more important than how many glycans are present when it comes to surface masking.

Circular dichroism (CD) spectroscopy shows no major structural changes among these glycoforms, so the observed differences must come from surface interactions, not overall protein folding^7^. Our simulations show that the N24 glycan mainly stays exposed to solvent, forming only weak and short-lived contacts with the protein. As such, N24 seems to act more like a hydrophilic decoration, adding general solubility and charge, but it does not actively shield the key hydrophobic helix near N83.

### Summary and conclusion

The detailed site-resolved analyses of EPO glycosylation converge on a central conclusion: glycan positioning—not merely occupancy or count—governs the molecular and pharmacological behavior of glycoproteins. This spatial encoding translates into distinct functional roles for the three N-glycosylation sites on EPO, which together form a tunable system for modulating structural exposure, folding efficiency, and surface hydropathy. Beyond EPO, this hierarchy may generalize to other glycoproteins with localized hydrophobic hotspots and multiple glycosylation sites. Our approach—integrating enhanced-sampling dynamics with site-specific experimental data—offers a scalable method to probe and optimize glycan functionality at atomic resolution. In doing so, it bridges structural biophysics with therapeutic design, enabling rational manipulation of glycosylation for improved drug developability, manufacturability, and in vivo performance.

In this work, we employed glycan-targeted enhanced-sampling molecular dynamics simulations to elucidate the molecular mechanisms by which site-specific N-glycosylation modulates the conformational landscape and surface properties of erythropoietin (EPO). Our findings demonstrate that N-glycans, particularly the glycan at Asn83, engage in persistent interactions with hydrophobic protein surfaces, leading to localized reductions in solvent-accessible hydropathy. This dynamic shielding effect correlates quantitatively with experimental retention times in hydrophobic interaction chromatography, providing a direct structure–property relationship. Moreover, principal component and RMSF analyses reveal that glycosylation induces long-range allosteric effects on distal loops and global protein flexibility. The observed cooperativity among glycosylation sites underscores the non-additive nature of glycan contributions to protein behavior. These results offer a mechanistic basis for the pharmacological advantages conferred by specific glycosylation patterns and support the rational design of glycoprotein therapeutics through modulation of glycan structure, occupancy, and topology.

## METHOD

### System Preparation

The glycoprotein structure of erythropoietin (EPO) was modeled using the Glycan Reader & Modeler module within CHARMM-GUI^17^. Based on the crystal structure of PDB ID 1BUY, amino acid mutations were introduced to replicate the recombinant EPO sequence, following the study by Murakami et al.^7^. N-linked glycans of the biantennary complex-type were attached at Asn24 (N24), Asn38 (N38), and Asn83 (N83). The topology and parameter files necessary for molecular simulations were generated with CHARMM-GUI using the CHARMM36m force field^35^. The solvated system was prepared by immersing the glycoprotein in a cubic periodic box filled with TIP3P water molecules, ensuring a minimum buffer distance of 15 Å from the protein surface. To neutralize the system charge and replicate physiological ionic conditions, Na⁺ and Cl⁻ ions were added to achieve a final ionic strength of 150 mM.

### Conventional Molecular Dynamics Simulations

All molecular dynamics simulations were carried out with the GENESIS simulation suite version 2.1^36–38^. Energy minimization was performed using the steepest descent method for 120,000 steps. Initially, positional restraints were applied to non-hydrogen atoms of the protein and glycans during the first 20,000 steps, focusing on minimizing the solvent and ion components. Subsequently, the entire system was subjected to an unrestrained minimization for the remaining 100,000 steps.

Equilibration was performed in three sequential phases. During the first phase, positional restraints were maintained on non-hydrogen glycoprotein atoms, while the system was gradually heated to 310 K over 200 ps in the NVT ensemble using the velocity Verlet integrator. In the second phase, the ensemble was switched to NPT conditions, and the system was equilibrated for an additional 200 ps at a pressure of 1 atm with the same positional restraints. For the third phase, all restraints were removed, and the equilibration process was repeated under identical NPT conditions. Long-range electrostatics were computed using the Particle Mesh Ewald (PME) algorithm^39, 40^, and van der Waals interactions were truncated at a 12 Å cutoff^41^. Temperature and pressure regulation was maintained using the stochastic velocity rescaling thermostat and MTK style barostat^42, 43^ with the refined temperature definition^44^. Group temperature/pressure is used for evaluation of temperature/pressure in thermostat/barostat^45, 46^.

For production runs, we utilized the r-RESPA integration with 3.5 and 7.0 fs time steps for the real- and reciprocal-space interactions, respectively^47^. Prior to the main simulations, further stabilization was ensured through two consecutive equilibration runs of 1.05 ns under NVT and NPT conditions, respectively, utilizing the multiple time step integrator. Subsequently, conventional molecular dynamics (cMD) simulations were performed in the NVT ensemble for a total of 525 ns, generating three independent trajectories per system. Simulation coordinates and energy profiles were saved every 10.5 ps throughout the production runs.

### Generalized Replica Exchange with Solute Tempering

We performed all-atom molecular dynamics (MD) simulations to investigate the conformational dynamics of erythropoietin (EPO) glycans. All combinations of N-linked glycosylation at Asn24, Asn38, and Asn83 (N24, N38, N83, N24N38, N24N83, N38N83, and N24N38N83) were simulated. To enhance sampling of glycan conformations attached to EPO, we applied generalized replica exchange with solute tempering (gREST), where only the glycan residues were defined as the solute region. In the gREST scheme, we selectively enhance the solute region by treating glycan internal torsional (dihedral) angles, Lennard-Jones interactions, and electrostatic interactions at higher temperatures than 310 K, while the rest of the system is simulated at room temperature (310 K). The number of replicas and the highest temperature was adjusted according to the number of attached glycans: 8 replicas and 545K for one glycan, 10 replicas and 516K for two glycans, and 12 replicas and 513K for three glycans. The temperature of each replica was automatically determined using the built-in optimization function in GENESIS, such that the exchange acceptance ratio between neighboring replicas was approximately 0.25. For each system, we first relaxed each replica at different solute temperatures for 10 ps followed by production runs for 525 ns per each replica. Replica-exchange attempts were performed every 10.5 ps. The full list of replica temperatures is provided in Table S1 of the Supporting Information.

### Structural Analysis

Root-mean-square fluctuation (RMSF) analysis was performed to quantify the structural flexibility of the EPO protein. All trajectories were first aligned to a reference structure based on the Cα atoms of the protein to remove overall translational and rotational motions. RMSF values were then calculated for each Cα atom. To characterize the spatial relationships between glycans and the protein, we defined the centers of mass (COM) for the protein, each glycan, and each glycosylation site (Asn24, Asn38, Asn83). Using these reference points, the following geometric metrics were computed: (1) Glycan–glycan distances: Euclidean distances between the COMs of each glycan pair. (2) Glycan–Asn–protein angles: The angle formed by three COMs—glycan, its attachment Asn residue, and the global protein COM. Distributions of these distances and angles were calculated over the simulation trajectories to evaluate the relative orientation and flexibility of glycans with respect to the protein scaffold. Principal component analysis (PCA)^32, 48^ was performed using the analysis tools provided by GENESIS. For each EPO construct, PCA was carried out separately for the protein and for the glycans, using Cartesian atomic coordinates. Prior to analysis, the protein structures were aligned to a reference conformation based on the Cα atoms of EPO to remove overall translational and rotational motions. PCA of the protein was then performed using the Cartesian coordinates of the Cα atoms. For the glycans, PCA was performed using the Cartesian coordinates of all heavy atoms (non-hydrogen atoms).

### Contact Analysis Between Glycans and the EPO Surface

The interactions between erythropoietin (EPO) and its conjugated N-glycans were quantitatively examined using structural snapshots sampled every 1.05 ns from gREST simulations. This analysis focused on characterizing glycan-protein contacts, particularly in the antenna regions of the glycans. A contact event was recorded when at least one atom from a glycan residue was found within 4.0 Å from any atom of an EPO residue. To prevent contact overestimation due to covalent linkages, the core N-acetylglucosamine residues, which are directly attached to the protein, were excluded from the evaluation. The analysis was thus directed towards the flexible antenna segments of the glycans, which play a more dynamic role in protein surface interactions. For each sampled structure, pairwise distances between atoms in the glycan antenna and the protein were computed to identify the closest atom-atom proximity. Contacts were accumulated across all trajectory frames, and contact frequencies were normalized by the total number of possible contact events, calculated as the product of the number of frames and the number of glycan antenna residues. This normalization allowed a residue-specific assessment of glycan interaction tendencies across the EPO surface.

### Calculation of Solvent-Accessible Surface Area (SASA)

SASA was calculated using CPPTRJ from the AmberTools suite^49^. To evaluate the exposure of hydrophobic regions in erythropoietin (EPO), SASA values were computed for both all residues and hydrophobic residues only, defined as Ala, Val, Leu, Ile, Met, Phe, Trp, and Pro. The calculations were performed on MD trajectories, and the resulting SASA values were averaged over time to obtain representative estimates of solvent accessibility. A probe radius of 1.4 Å was used to represent the size of a water molecule. SASA was computed for the following residue ranges to examine local and global surface properties: (1) Residues 55–85, (2) Residues 55–115, (3) Residues 55–85 and 115–130 (combined), (4) The entire protein (residues 1–166).

## Acknowledgments

This work was supported in part by MEXT JSPS KAKENHI [Grant Nos. 21H05249 (to Y.S.)]; RIKEN pioneering projects “Biology of Intracellular Environments” (to Y.S.); MEXT program for Big-data-driven bio/synthetic polymer science to create absolutely circular materials (JPMXP1122714694) and Data-Driven Research Methods Development and Materials Innovation Led by Computational Materials Science (JPMXP1020230327) (to Y.S.). The authors thank Yasuhiro Kajihara for helpful discussions on this work.

## References

(1) Schjoldager, K. T.; Narimatsu, Y.; Joshi, H. J.; Clausen, H. Global view of human protein glycosylation pathways and functions. Nature Reviews Molecular Cell Biology 2020, 21 (12), 729–749. DOI: 10.1038/s41580-020-00294-x.

(2) Moremen, K. W.; Tiemeyer, M.; Nairn, A. V. Vertebrate protein glycosylation: diversity, synthesis and function. Nature Reviews Molecular Cell Biology 2012, 13 (7), 448–462. DOI: 10.1038/nrm3383.

(3) Gazaway, E.; Kandel, R.; Grant, O. C.; Woods, R. J. Are N-linked glycans intrinsically disordered? Current Opinion in Structural Biology 2025, 93, 103118. DOI: 10.1016/j.sbi.2025.103118.

(4) Kato, K.; Yanaka, S.; Yamaguchi, T. The synergy of experimental and computational approaches for visualizing glycoprotein dynamics: Exploring order within the apparent disorder of glycan conformational ensembles. Current Opinion in Structural Biology 2025, 92, 103049. DOI: 10.1016/j.sbi.2025.103049.

(5) Peng, B.; Kong, G. C.; Yang, C.; Ming, Y. Z. Erythropoietin and its derivatives: from tissue protection to immune regulation. Cell Death Dis 2020, 11 (2). DOI: 10.1038/s41419-020-2276-8.

(6) Suresh, S.; Rajvanshi, P. K.; Noguchi, C. T. The Many Facets of Erythropoietin Physiologic and Metabolic Response. Front Physiol 2020, 10. DOI: ARTN 1534 10.3389/fphys.2019.01534.

(7) Murakami, M.; Kiuchi, T.; Nishihara, M.; Tezuka, K.; Okamoto, R.; Izumi, M.; Kajihara, Y. Chemical synthesis of erythropoietin glycoforms for insights into the relationship between glycosylation pattern and bioactivity. Sci Adv 2016, 2 (1). DOI: ARTN e1500678 10.1126/sciadv.1500678.

(8) Li, C.; Wang, L. X. Chemoenzymatic Methods for the Synthesis of Glycoproteins. Chem Rev 2018, 118 (17), 8359–8413. DOI: 10.1021/acs.chemrev.8b00238.

(9) Nomura, K.; Maki, Y.; Okamoto, R.; Satoh, A.; Kajihara, Y. Glycoprotein Semisynthesis by Chemical Insertion of Glycosyl Asparagine Using a Bifunctional Thioacid-Mediated Strategy. J Am Chem Soc 2021, 143 (27), 10157–10167. DOI: 10.1021/jacs.1c02601.

(10) Dong, W.; Yang, X.; Li, X.; Wei, S.; An, C.; Zhang, J.; Shi, X.; Dong, S. Investigation of N-Glycan Functions in Receptor for Advanced Glycation End Products V Domain through Chemical Glycoprotein Synthesis. J Am Chem Soc 2024, 146 (27), 18270–18280. DOI: 10.1021/jacs.4c01413.

(11) Kiuchi, T.; Izumi, M.; Mukogawa, Y.; Shimada, A.; Okamoto, R.; Seko, A.; Sakono, M.; Takeda, Y.; Ito, Y.; Kajihara, Y. Monitoring of Glycoprotein Quality Control System with a Series of Chemically Synthesized Homogeneous Native and Misfolded Glycoproteins. J Am Chem Soc 2018, 140 (50), 17499–17507. DOI: 10.1021/jacs.8b08653.

(12) Li, H. X.; Zhang, J.; An, C. J.; Dong, S. W. Probing -Glycan Functions in Human Interleukin-17A Based on Chemically Synthesized Homogeneous Glycoforms. J Am Chem Soc 2021, 143 (7), 2846–2856. DOI: 10.1021/jacs.0c12448.

(13) Liu, Y. B.; Nomura, K.; Abe, J.; Kajihara, Y. Recent advances on the synthesis of N-linked glycoprotein for the elucidation of glycan functions. Curr Opin Chem Biol 2023, 73. DOI: ARTN 102263 10.1016/j.cbpa.2023.102263.

(14) Maki, Y.; Okamoto, R.; Izumi, M.; Kajihara, Y. Chemical Synthesis of an Erythropoietin Glycoform Having a Triantennary N-Glycan: Significant Change of Biological Activity of Glycoprotein by Addition of a Small Molecular Weight Trisaccharide. J Am Chem Soc 2020, 142 (49), 20671–20679. DOI: 10.1021/jacs.0c08719 From NLM Medline.

(15) Abramson, J.; Adler, J.; Dunger, J.; Evans, R.; Green, T.; Pritzel, A.; Ronneberger, O.; Willmore, L.; Ballard, A. J.; Bambrick, J.;, et al. Accurate structure prediction of biomolecular interactions with AlphaFold 3. Nature 2024, 630 (8016). DOI: 10.1038/s41586-024-07487-w.

(16) Jumper, J.; Evans, R.; Pritzel, A.; Green, T.; Figurnov, M.; Ronneberger, O.; Tunyasuvunakool, K.; Bates, R.; Zídek, A.; Potapenko, A.;, et al. Highly accurate protein structure prediction with AlphaFold. Nature 2021, 596 (7873), 583-+. DOI: 10.1038/s41586-021-03819-2.

(17) Park, S. J.; Lee, J.; Qi, Y. F.; Kern, N. R.; Lee, H. S.; Jo, S.; Joung, I.; Joo, K.; Lee, J.; Im, W. CHARMM-GUI Glycan Modeler for modeling and simulation of carbohydrates and glycoconjugates. Glycobiology 2019, 29 (4), 320–331. DOI: 10.1093/glycob/cwz003.

(18) Yang, M. J.; Huang, J.; MacKerell, A. D. Enhanced Conformational Sampling Using Replica Exchange with Concurrent Solute Scaling and Hamiltonian Biasing Realized in One Dimension. J Chem Theory Comput 2015, 11 (6), 2855–2867. DOI: 10.1021/acs.jctc.5b00243.

(19) Mori, T.; Jung, J.; Kobayashi, C.; Dokainish, H. M.; Re, S.; Sugita, Y. Elucidation of interactions regulating conformational stability and dynamics of SARS-CoV-2 S-protein. Biophys J 2021, 120 (6), 1060–1071. DOI: 10.1016/j.bpj.2021.01.012.

(20) Sztain, T.; Ahn, S. H.; Bogetti, A. T.; Casalino, L.; Goldsmith, J. A.; Seitz, E.; McCool, R. S.; Kearns, F. L.; Acosta-Reyes, F.; Maji, S.;, et al. A glycan gate controls opening of the SARS-CoV-2 spike protein. Nat Chem 2021, 13 (10), 963-+. DOI: 10.1038/s41557-021-00758-3.

(21) Dokainish, H. M.; Re, S.; Mori, T.; Kobayashi, C.; Jung, J. W.; Sugita, Y. The inherent flexibility of receptor binding domains in SARS-CoV-2 spike protein. Elife 2022, 11. DOI: ARTN e75720 10.7554/eLife.75720.

(22) Dokainish, H. M.; Sugita, Y. Structural effects of spike protein D614G mutation in SARS-CoV-2. Biophys J 2023, 122 (14), 2910–2920. DOI: 10.1016/j.bpj.2022.11.025.

(23) Sugita, Y.; Okamoto, Y. Replica-exchange molecular dynamics method for protein folding. Chem Phys Lett 1999, 314 (1-2), 141–151. DOI: Doi 10.1016/S0009-2614(99)01123-9.

(24) Re, S.; Miyashita, N.; Yamaguchi, Y.; Sugita, Y. Structural Diversity and Changes in Conformational Equilibria of Biantennary Complex-Type -Glycans in Water Revealed by Replica-Exchange Molecular Dynamics Simulation. Biophys J 2011, 101 (10), L44–L46. DOI: 10.1016/j.bpj.2011.10.019.

(25) Sugita, Y.; Nishima, W.; Yamaguchi, Y.; Re, S. Y. Conformational Diversity of N-Glycans in Solution Studied by REMD Simulations. Biophys J 2013, 104 (2), 170a–170a. DOI: DOI 10.1016/j.bpj.2012.11.959.

(26) Galvelis, R.; Re, S. Y.; Sugita, Y. Enhanced Conformational Sampling of N-Glycans in Solution with Replica State Exchange Metadynamics. J Chem Theory Comput 2017, 13 (5), 1934–1942. DOI: 10.1021/acs.jctc.7b00079.

(27) Fukunishi, H.; Watanabe, O.; Takada, S. On the Hamiltonian replica exchange method for efficient sampling of biomolecular systems: Application to protein structure prediction. J Chem Phys 2002, 116 (20), 9058–9067. DOI: 10.1063/1.1472510.

(28) Mishra, S. K.; Kara, M.; Zacharias, M.; Koca, J. Enhanced conformational sampling of carbohydrates by Hamiltonian replica-exchange simulation. Glycobiology 2014, 24 (1), 70–84. DOI: 10.1093/glycob/cwt093.

(29) Gil-Ley, A.; Bussi, G. Enhanced Conformational Sampling Using Replica Exchange with Collective-Variable Tempering (vol 11, pg 1077, 2015). J Chem Theory Comput 2015, 11 (11), 5554–5554. DOI: 10.1021/acs.jctc.5b00981.

(30) Grothaus, I. L.; Bussi, G.; Ciacchi, L. C. Exploration, Representation, and Rationalization of the Conformational Phase Space of N-Glycans. J Chem Inf Model 2022, 62 (20), 4992–5008. DOI: 10.1021/acs.jcim.2c01049.

(31) Kamiya, M.; Sugita, Y. Flexible selection of the solute region in replica exchange with solute tempering: Application to protein-folding simulations. J Chem Phys 2018, 149 (7). DOI: Artn 072304 10.1063/1.5016222.

(32) Kitao, A.; Hirata, F.; Go, N. The Effects of Solvent on the Conformation and the Collective Motions of Protein - Normal Mode Analysis and Molecular-Dynamics Simulations of Melittin in Water and in Vacuum. Chem Phys 1991, 158 (2-3), 447–472. DOI: Doi 10.1016/0301-0104(91)87082-7.

(33) Ives, C. M.; Singh, O.; D’Andrea, S.; Fogarty, C. A.; Harbison, A. M.; Satheesan, A.; Tropea, B.; Fadda, E. Restoring protein glycosylation with GlycoShape. Nat Methods 2024, 21 (11). DOI: 10.1038/s41592-024-02464-7.

(34) Tsai, Y. X.; Chang, N. E.; Reuter, K.; Chang, H. T.; Yang, T. J.; von Bulow, S.; Sehrawat, V.; Zerrouki, N.; Tuffery, M.; Gecht, M.;, et al. Rapid simulation of glycoprotein structures by grafting and steric exclusion of glycan conformer libraries. Cell 2024, 187 (5). DOI: 10.1016/j.cell.2024.01.034.

(35) Huang, J.; Rauscher, S.; Nawrocki, G.; Ran, T.; Feig, M.; de Groot, B. L.; Grubmüller, H.; MacKerell, A. D. CHARMM36m: an improved force field for folded and intrinsically disordered proteins. Nat Methods 2017, 14 (1), 71–73. DOI: 10.1038/Nmeth.4067.

(36) Jung, J.; Mori, T.; Kobayashi, C.; Matsunaga, Y.; Yoda, T.; Feig, M.; Sugita, Y. GENESIS: a hybrid-parallel and multi-scale molecular dynamics simulator with enhanced sampling algorithms for biomolecular and cellular simulations. Wires Comput Mol Sci 2015, 5 (4), 310–323. DOI: 10.1002/wcms.1220.

(37) Kobayashi, C.; Jung, J.; Matsunaga, Y.; Mori, T.; Ando, T.; Tamura, K.; Kamiya, M.; Sugita, Y. GENESIS 1.1: A hybrid-parallel molecular dynamics simulator with enhanced sampling algorithms on multiple computational platforms. J Comput Chem 2017, 38 (25), 2193–2206. DOI: 10.1002/jcc.24874.

(38) Jung, J.; Yagi, K.; Tan, C.; Oshima, H.; Mori, T.; Yu, I.; Matsunaga, Y.; Kobayashi, C.; Ito, S.; La Torre, D. U.;, et al. GENESIS 2.1: High-Performance Molecular Dynamics Software for Enhanced Sampling and Free-Energy Calculations for Atomistic, Coarse-Grained, and Quantum Mechanics/Molecular Mechanics Models. J Phys Chem B 2024, 128 (25), 6028–6048. DOI: 10.1021/acs.jpcb.4c02096.

(39) Darden, T.; York, D.; Pedersen, L. Particle Mesh Ewald - an N.Log(N) Method for Ewald Sums in Large Systems. J Chem Phys 1993, 98 (12), 10089–10092. DOI: Doi 10.1063/1.464397.

(40) Essmann, U.; Perera, L.; Berkowitz, M. L.; Darden, T.; Lee, H.; Pedersen, L. G. A Smooth Particle Mesh Ewald Method. J Chem Phys 1995, 103 (19), 8577–8593. DOI: Doi 10.1063/1.470117.

(41) Steinbach, P. J.; Brooks, B. R. New Spherical-Cutoff Methods for Long-Range Forces in Macromolecular Simulation. J Comput Chem 1994, 15 (7), 667–683. DOI: DOI 10.1002/jcc.540150702.

(42) Bussi, G.; Donadio, D.; Parrinello, M. Canonical sampling through velocity rescaling. J Chem Phys 2007, 126 (1). DOI: Artn 014101 10.1063/1.2408420.

(43) Martyna, G. J.; Tuckerman, M. E.; Tobias, D. J.; Klein, M. L. Explicit reversible integrators for extended systems dynamics. Mol Phys 1996, 87 (5), 1117–1157. DOI: Doi 10.1080/00268979600100761.

(44) Jung, J.; Kobayashi, C.; Sugita, Y. Optimal Temperature Evaluation in Molecular Dynamics Simulations with a Large Time Step. J Chem Theory Comput 2019, 15 (1), 84–94. DOI: 10.1021/acs.jctc.8b00874.

(45) Lippert, R. A.; Predescu, C.; Ierardi, D. J.; Mackenzie, K. M.; Eastwood, M. P.; Dror, R. O.; Shaw, D. E. Accurate and efficient integration for molecular dynamics simulations at constant temperature and pressure. J Chem Phys 2013, 139 (16). DOI: Artn 164106 10.1063/1.4825247.

(46) Jung, J.; Sugita, Y. Group-based evaluation of temperature and pressure for molecular dynamics simulation with a large time step. J Chem Phys 2020, 153 (23). DOI: Artn 234115 10.1063/5.0027873.

(47) Tuckerman, M.; Berne, B. J.; Martyna, G. J. Reversible Multiple Time Scale Molecular-Dynamics. J Chem Phys 1992, 97 (3), 1990–2001. DOI: Doi 10.1063/1.463137.

(48) Jolliffe, I. Principal Component Analysis. In International Encyclopedia of Statistical Science, Lovric, M. Ed.; Springer Berlin Heidelberg, 2011; pp 1094–1096.

(49) Roe, D. R.; Cheatham, T. E., III. PTRAJ and CPPTRAJ: Software for Processing and Analysis of Molecular Dynamics Trajectory Data. J Chem Theory Comput 2013, 9 (7), 3084–3095. DOI: 10.1021/ct400341p.

